# Gene Silencing in Plants by Artificial Small RNAs Derived from Minimal Precursors and Expressed via Tobacco Rattle Virus

**DOI:** 10.1101/2025.03.27.645721

**Authors:** María Juárez-Molina, Ana Alarcia, Anamarija Primc, Iván Ortega-Miralles, Adriana E. Cisneros, Alberto Carbonell

## Abstract

Highly specific, second-generation RNA interference tools are based on artificial small RNAs (art-sRNAs), such as artificial microRNAs (amiRNAs) and synthetic trans-acting small interfering RNAs (syn-tasiRNAs). Recent progress includes the use of minimal-length precursors to express art-sRNAs in plants. These minimal precursors retain the minimal structural elements for recognition and efficient processing by host enzymes. They yield high amounts of art-sRNAs and remain stable when incorporated into potato virus X-based viral vectors for art-sRNA-mediated virus-induced gene silencing (art-sRNA-VIGS). However, further adaptation to new viral vector systems with reduced symptomatology is needed to improve the versatility of art-sRNA-VIGS. Here, we developed a novel platform based on tobacco rattle virus (TRV) –a widely used viral vector inducing minimal or no symptoms– for the delivery of art-sRNAs into plants. TRV was engineered to express authentic amiRNAs and syn-tasiRNAs from minimal precursors in *Nicotiana benthamiana*, resulting in robust and highly specific silencing of endogenous genes. Notably, the expression of syn-tasiRNAs through TRV conferred strong resistance against tomato spotted wilt virus, an economically important pathogen. Furthermore, we established a transgene-free approach by applying TRV-containing crude extracts through foliar spraying, eliminating the need for stable genetic transformation. In summary, our results highlight the unique advantages of minimal precursors and extend the application of art-sRNA-VIGS beyond previously established viral vector systems, providing a scalable, rapid and highly specific tool for gene silencing.

**Key message:** We developed a novel tobacco rattle virus-based platform for the transgene-free expression of both artificial microRNAs and synthetic trans-acting small interfering RNAs for efficient gene silencing in plants.

## INTRODUCTION

Gene silencing by small RNAs (sRNAs) is a fundamental regulatory mechanism in plants that controls gene expression at transcriptional and post-transcriptional levels (Axtell 2013; Bologna and Voinnet 2014). Classic gene silencing approaches based on long double-stranded RNAs (dsRNAs), such as virus-induced gene silencing (VIGS) and hairpin RNA (hpRNA)-mediated silencing, have limited specificity. These methods generate large pools of small interfering RNAs (siRNAs), some of which can accidentally target cellular RNAs with sequence complementarity and lead to off-target effects and potential toxicity (Jackson et al. 2003). To overcome this limitation, highly specific second-generation gene silencing tools based on artificial small RNAs (art-sRNAs) have been developed as efficient alternatives for targeted gene silencing in plants. Art-sRNAs are 21-nucleotide (nt) sRNAs designed computationally to bind and cleave target RNAs with high efficiency and specificity, and with no off-target effects (Carbonell 2017). The two main classes of art-sRNAs are artificial microRNAs (amiRNAs) and synthetic trans-acting small interfering RNAs (syn-tasiRNAs), which are functionally similar but differ in their biogenesis pathway. AmiRNAs are derived from modified endogenous miRNA precursors, where the native miRNA/miRNA* duplex is replaced by a designed amiRNA/amiRNA* sequence (Schwab et al. 2006). These precursors are processed by DICER-LIKE1 (DCL1) into mature amiRNAs, which are then incorporated into ARGONAUTE1 (AGO1) to direct target RNA cleavage. Syn-tasiRNAs, on the other hand, originate from *TAS* precursors, which are first cleaved by a specific miRNA-AGO complex (de la Luz Gutierrez-Nava et al. 2008; Zhang 2014). One cleavage product is stabilized and converted into dsRNA by RNA-DEPENDENT RNA POLYMERASE6 (RDR6) (Allen et al. 2005; Yoshikawa et al. 2005), followed by sequential processing by DCL4 into phased 21-nt syn-tasiRNAs, which associate with AGO1 to silence target transcripts. Importantly, while amiRNAs are typically designed to silence individual genes, syn-tasiRNAs allow for multiplex targeting by producing multiple art-sRNAs from a single precursor, making them particularly effective in achieving multi-gene silencing and durable antiviral protection (Carbonell 2019; Cisneros and Carbonell 2020).

Despite their versatility, the application of art-sRNAs in plants has been constrained by the requirement to transgenically express long precursor transcripts. This limitation has been recently addressed by the engineering of minimal precursors, which retain the essential structural features required for accurate processing while significantly reducing their overall length. For instance, the *shc* minimal amiRNA precursor is only 89-nt long, and includes the *AtMIR390a* basal stem, the amiRNA/amiRNA* duplex, and a deleted version of the *OsMIR390* distal stem-loop (Cisneros et al. 2023). On the other hand, minimal syn-tasiRNA precursors consist of a 22-nt endogenous miRNA target site (TS) followed by an 11-nt spacer and the 21-n syn-tasiRNA sequence(s) (Cisneros et al. 2025). Remarkably, minimal but not full-length art-sRNA precursors produced authentic amiRNAs or syn-tasiRNAs and induced widespread gene silencing in *N. benthamiana* when expressed from an RNA virus such as potato virus X (PVX), which can be applied by spraying infectious crude extracts onto leaves in a GMO-free manner (Cisneros et al. 2023; Cisneros et al. 2025). These strategies, named amiRNA-based VIGS (amiR-VIGS) or syn-tasiRNA-based VIGS (syn-tasiR-VIGS), were further used to vaccinate plants against a pathogenic virus, resulting in complete plant immunization in the case of syn-tasiR-VIGS (Cisneros et al. 2025). Still, PVX-based art-sRNA-VIGS in *N. benthamiana* induce mild symptoms that may interfere with the expected gene silencing phenotypes, at least in particular cases. In addition, to date, art-sRNA-VIGS using minimal precursors has only been reported with PVX. Therefore, art-sRNA-VIGS requires further adaptation to new viral vector systems, particularly those limiting viral vector symptomatology, for broadening and optimizing its applicability.

Here, we present a tobacco rattle virus (TRV)-based platform for producing amiRNAs and syn-tasiRNAs in plants for highly efficient and widespread gene silencing. We show that authentic amiRNA and syn-tasiRNAs can be produced in *N. benthamiana* through TRV-based amiR-VIGS and syn-tasiR-VIGS, respectively, for silencing endogenous genes with minimal or no TRV-derived symptoms. Moreover, TRV-based syn-tasiR-VIGS induced high antiviral resistance against the economically important tomato spotted wilt virus (TSWV) plant pathogen. Importantly, we established the transgene-free delivery of TRV-based art-sRNA-VIGS to plants by spraying crude extracts, thus allowing for widespread gene silencing without the need for genetic transformation. Our findings extend the use of art-sRNA-VIGS to a new viral vector system and highlight the potential of TRV-based art-sRNA expression from minimal precursors as a scalable and efficient tool for functional genomics and crop protection.

## MATERIALS AND METHODS

### Plant species and growth conditions

*N. benthamiana* plants were cultivated in a growth chamber set at 25°C under a 12-hour light/12-hour-dark photoperiod. Plant images were captured using a Nikon D3000 digital camera equipped with an AF-S DX NIKKOR 18-55 mm f/3.5–5.6G VR lens.

### Artificial small RNA design

AmiR-NbSu, syn-tasiR-NbSu, syn-tasiR-GUS_Sl_-1, syn-tasiR-GUS_Sl_-2, syn-tasiR-TSWV-1, syn-tasiR-TSWV-2, syn-tasiR-TSWV-3 and syn-tasiR-TSWV-4 guide sequences were described before (Carbonell et al. 2019; Cisneros et al. 2022).

### DNA constructs

For TRV-based amiRNA constructs, amiRNA cassettes pri-amiR-NbSu and shc-amiR-NbSu were amplified from *35S:AtMIR390a-NbSu-2* (Addgene plasmid #213400) (Cisneros et al. 2022) with oligonucleotide pair AC-615/AC-616 and AC-617/AC-618 respectively, and gel purified. For TRV-based syn-tasiRNA constructs, syn-tasiRNA cassettes *TAS1c-miR482TS-Su*, *miR173TS-Su*, *miR482TS-TSWV(x4)* and miR173TS-TSWV(x4) were amplified from *35S:AtTAS1c(NbmiR482aTS)-D2-NbSu*, *35S:PVX-min_173_-Su*, *35S:PVXmin_482_-TSWV(x4)* and *35S:PVXmin_173_-TSWV(x4)* (Cisneros et al. 2025) with oligonucleotide pairs AC-518/AC-519, AC-1222/AC-1223, AC-985/AC-986 and AC-1222/AC-986, respectively, and gel purified. Syn-tasiRNA cassettes *miR482TS-Su* and *miR482TS-GUS(x4)* were ordered as dsDNA oligonucleotides AC-667 and AC-984, respectively. All amiRNA and syn-tasiRNA cassettes were assembled into *Bsa*I-digested and gel-purified *pLX-TRV2* (Addgene plasmid #180516) (Aragonés et al. 2022) in the presence of GeneArt Gibson Assembly HiFi Master Mix (Invitrogen) to generate *35S:TRV2-pri-amiR-Su*, *35S:TRV2-shc-amiR-Su*, *35S:TRV2-TAS1c-miR482TS-Su*, *35S:TRV2-miR482TS-Su*, *35S:TRV2-miR173TS-Su, 35S:TRV2-miR482TS-GUS(x4)* and *35S:TRV2-miR482TS-TSWV(x4)*. *35S:TRV1* and *35S:TRV2* were described before (Aragonés et al. 2022). A detailed protocol for cloning amiRNAs or syn-tasiRNA minimal precursors into *pLB-TRV2* is described in Text S1. The sequences of all syn-tasiRNA precursors are listed in Text S2.

### Transient expression of constructs and spray-based inoculation of viruses

Agrobacterium-mediated infiltration of DNA constructs into *N. benthamiana* leaves was performed as described previously (Llave et al. 2002; Cuperus et al. 2010). The preparation and spraying of crude extracts derived from virus-infected *N. benthamiana* plants followed established protocols (Cisneros et al. 2023), with 5% silicon carbide (carborundum) included in the inoculation buffer.

### RNA preparation

Total RNA was extracted from *N. benthamiana* leaves as previously described (Cisneros et al. 2023). Triplicate samples were prepared from pools consisting of two apical leaves.

### Real-time RT-qPCR

cDNA was synthesized from 500 ng of DNase I-treated total RNA extracted from *N. benthamiana* leaves using the PrimeScript RT Reagent Kit (Perfect Real Time, Takara), according to the manufacturer’s instructions. Real-time RT-qPCR was performed using the same RNA samples previously used for sRNA blot analysis as described (Cisneros et al. 2025). Oligonucleotides used for RT-qPCR are listed in Table S1. Target mRNA expression levels were normalized to the reference genes *PP2A*, and relative expression was calculated using the delta-delta Ct method via QuantStudio Design and Analysis Software version 1.5.1 (Thermo Fisher Scientific). Three independent biological replicates were analyzed, each with two technical replicates.

### Stability and sequence analyses of syn-tasiRNA precursors during viral infections

Total RNA from the apical leaves of three biological replicates was pooled prior to cDNA synthesis. PCR was conducted to detect amiRNA or syn-tasiRNA precursors, TRV, and *PP2A* using the oligonucleotide pairs AC-523/AC-524, AC-660/AC-661, and AC-365/AC-366, respectively (Table S1), along with Phusion DNA Polymerase (Thermo Fisher Scientific). PCR products were analyzed through agarose gel electrophoresis, and bands of the expected size were excised and sequenced as needed.

### Small RNA blot assays

Small RNA blot assays and band quantification from radioactive membranes were performed as previously described (Cisneros et al. 2022). The oligonucleotides used as probes for sRNA blots are detailed in Table S1.

### Small RNA sequencing and data analysis

The quantity, purity, and integrity of total RNA were evaluated using a 2100 Bioanalyzer (RNA 6000 Nano kit, Agilent) before submission to BGI (Hong Kong, China) for sRNA library construction and SE50 high-throughput sequencing on a DNBSEQ-G-400 sequencer. Quality-trimmed and adaptor-removed clean reads provided by BGI were processed using the *fastx_collapser* toolkit (http://hannonlab.cshl.edu/fastx_toolkit) (Hannon 2010) to collapse identical reads into unique sequences while retaining read counts. Clean, unique reads were mapped to the forward strand of the syn-tasiRNA precursor expressed in each sample (Data S1) using a custom Python script, which allowed no mismatches or gaps, and calculated read counts and RPMs (reads per million mapped reads) for each mapping position.

The processing accuracy of syn-tasiRNA precursors was evaluated by quantifying the proportion of 19–24 nt sRNA (+) reads mapping within ± 4 nt of the 5′ end of the syn-tasiRNA guide, as described previously (Cuperus et al. 2010; Carbonell et al. 2015). Phasing register tables were generated by calculating the proportion of 21-nt sRNA (+) reads in each register relative to the corresponding sRNA cleavage site for all 21-nt positions downstream of the cleavage site, as before (Carbonell et al. 2014).

### Protein blot analysis

Proteins were resolved on NuPAGE Novex 4–12% Bis-Tris gels (Invitrogen), transferred to Protran nitrocellulose membranes (Amersham), and detected using specific antibodies with chemiluminescence as described (Cisneros et al. 2025).

### Gene and virus identifiers

*N. benthamiana* gene identifiers are *Su* (Nbv5.1tr6204879) and *PP2A* (Nbv5.1tr6224808), TSWV LL-N.05 segment L, M and S genome identifiers are KP008128, FM163373 and KP008129, respectively. *Escherichia coli* b-glucuronidase gene sequence corresponds to GenBank accession number S69414.1.

## RESULTS

### Gene silencing by amiRNAs derived from minimal precursors and expressed from TRV

The use of minimal amiRNA precursors for stable expression from viral vectors and effective silencing of plant genes has been recently reported employing the PVX vector (Cisneros et al. 2023). Given the limited cargo capacity of viral vectors, we hypothesized that minimal amiRNA precursors might be more effective than full-length precursors in other VIGS systems, such as those based on TRV. To test this, we designed the *pri-amiR-Su* and *shc-amiR-Su* precursors to express an amiRNA targeting the *N. benthamiana* magnesium chelatase subunit CHLI-encoding *SULPHUR* (*Su*) gene (Figure 1A). Silencing of *Su* induces a bleaching phenotype in affected tissues (Cisneros et al. 2022; Cisneros and Carbonell 2025). These precursors were derived from full-length *Arabidopsis thaliana* (Arabidopsis) *MIR390a* “*pri*” or minimal “*shc*” versions and were inserted into a TRV infectious clone containing TRV genome 2 (TRV2) to generate the *35S:TRV2-pri-amiR-Su* and *35S:TRV2-shc-amiR-Su* constructs, respectively (Figure 1B). If functional amiR-Su was produced, TRV-infected tissues were expected to bleach. These constructs, along with an insert-free *35S:TRV2* control construct, were independently agroinoculated into a single leaf of three *N. benthamiana* plants. A “mock” group of plants was agroinfiltrated with the agroinfiltration solution alone. To trigger TRV infections, TRV2-based constructs were transformed in an *Agrobacterium tumefaciens* C58C1 strain carrying the *pLX-TRV1* plasmid containing TRV genome 1.

**Figure 1.**
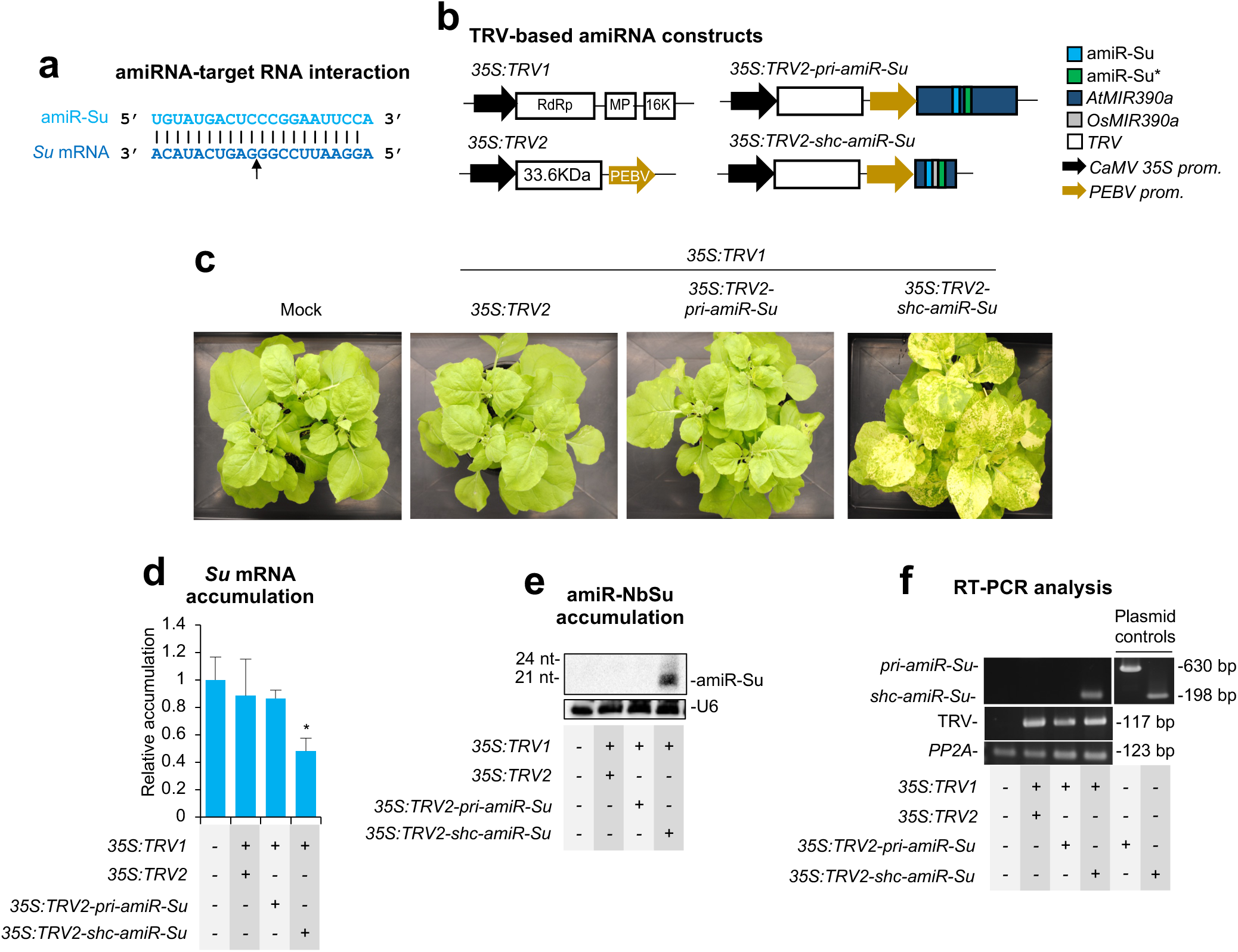
Functional analysis of tobacco rattle virus (TRV) constructs expressing amiR-Su from full-length (*pri*) *AtMIR390a* or minimal *shc*-based amiRNA precursors in *Nicotiana benthamiana*. (**a**) Base-pairing between amiR-Su and *Su* target mRNA. Nucleotides corresponding to the guide strand of the amiRNA or to the target mRNA are in light and dark blue, respectively. The arrow indicates the amiRNA-predicted cleavage site. (**b**) Diagram of TRV-based constructs. *AtMIR390a*, *OsMIR390*, amiR-Su and amiR-Su* sequences are represented by dark blue, grey, light blue and green and boxes, respectively. TRV ORFs and *35S*-based promoters are represented as white boxes and black arrows, respectively. RdRP, RNA-dependent RNA-polymerase; MP, movement protein; 16K, 16KDa protein; 33.6K, 33.6KDa protein; PEBV, pea early browning virus coat protein promoter. (**c**) Photos at 14 days post-agroinoculation (dpa) of sets of three plants agroinoculated with the different constructs. (**d**) Target *Su* mRNA accumulation in RNA preparations from apical leaves collected at 7 dpa and analysed individually (mock = 1.0 in all comparisons). Bars with an asterisk indicate whether the mean values are significantly different from mock control samples (*P* < 0.05 in pairwise Student’s *t*-test comparison). (**e**) Northern blot detection of amiR-Su in RNA preparations from apical leaves collected at 7 dpa and pooled from three independent plants. (**f**) RT-PCR detection of TRV and amiRNA precursors in apical leaves at 7 dpa. RT-PCR products corresponding to the *PP2A* are also shown as control (bottom), as well as positive control amplifications of *pri* and *shc* fragments from plasmids.

Bleaching of apical leaves was first observed at 8-9 days post-agroinoculation (dpa), but only in plants expressing *35S:TRV2-shc-amiR-Su*. By 14 dpa, bleaching had extended to most apical leaves (Figure 1C). At this time point, no TRV-derived symptoms were observed in control plants expressing *35S:TRV2*, which were phenotypically indistinguishable from mock-inoculated plants (Figure 1C), as observed before (Ratcliff et al. 2001). Plants were monitored until 28 dpa, and only those expressing *35S:TRV2-shc-amiR-Su* exhibited sustained *Su* silencing-associated bleaching. RT-qPCR and RNA-blot analyses confirmed that only plants expressing *35S:TRV2-shc-amiR-Su* accumulated low levels of *Su* mRNA (Figure 1D) and high levels of amiR-Su (Figure 1E). Additionally, RT-PCR analysis at 14 dpa revealed the presence of the minimal *shc-amiR-Su* precursor, while the full-length *pri-amiR-Su* precursor was not detected (Figure 1F). Since TRV was detected in plants expressing each of the precursors, the failure to detect the full-length *pri-amiR-Su* precursor likely reflects its deletion during TRV replication. These results indicate that minimal but not full-length precursors allow efficient amiRNA production from TRV and widespread gene silencing in *N. benthamiana*.

### Gene silencing by syn-tasiRNAs derived from minimal precursors and expressed from TRV

We next examined whether the TRV-based VIGS system could also support syn-tasiRNA production in *N. benthamiana*. Based on recent studies using PVX-based syn-tasiR-VIGS (Cisneros et al. 2025), we hypothesized that a minimal syn-tasiRNA precursor, such as *miR482TS*, containing *N. benthamiana* miR482 target site (TS) followed by an 11-nt spacer derived from Arabidopsis *TAS1c* (Figure 2A), would offer an advantage over full-length *TAS1c*-based precursors, due to its shorter length which should improve stability in the viral genome. To test this, *TAS1c-miR482TS-Su* and *miR482aTS-Su* sequences –engineered to express a syn-tasiRNA against *Su* (syn-tasiR-Su) (Figure 2A), which shares the same sequence as amiR-Su– from full-length *TAS1c* or minimal miR482TS-based precursors, respectively, were inserted into TRV2 to generate the *35S:TRV2-TAS1c-miR482TS-Su* and *35S:TRV2-miR482TS-Su* constructs (Figure 2B). Each construct was agroinoculated into a single leaf of three *N. benthamiana* plants, alongside an insert-free *35S:TRV2* control and a mock group. The bleaching phenotype (indicative of *Su* silencing) was monitored during 28 dpa, as before.

**Figure 2.**
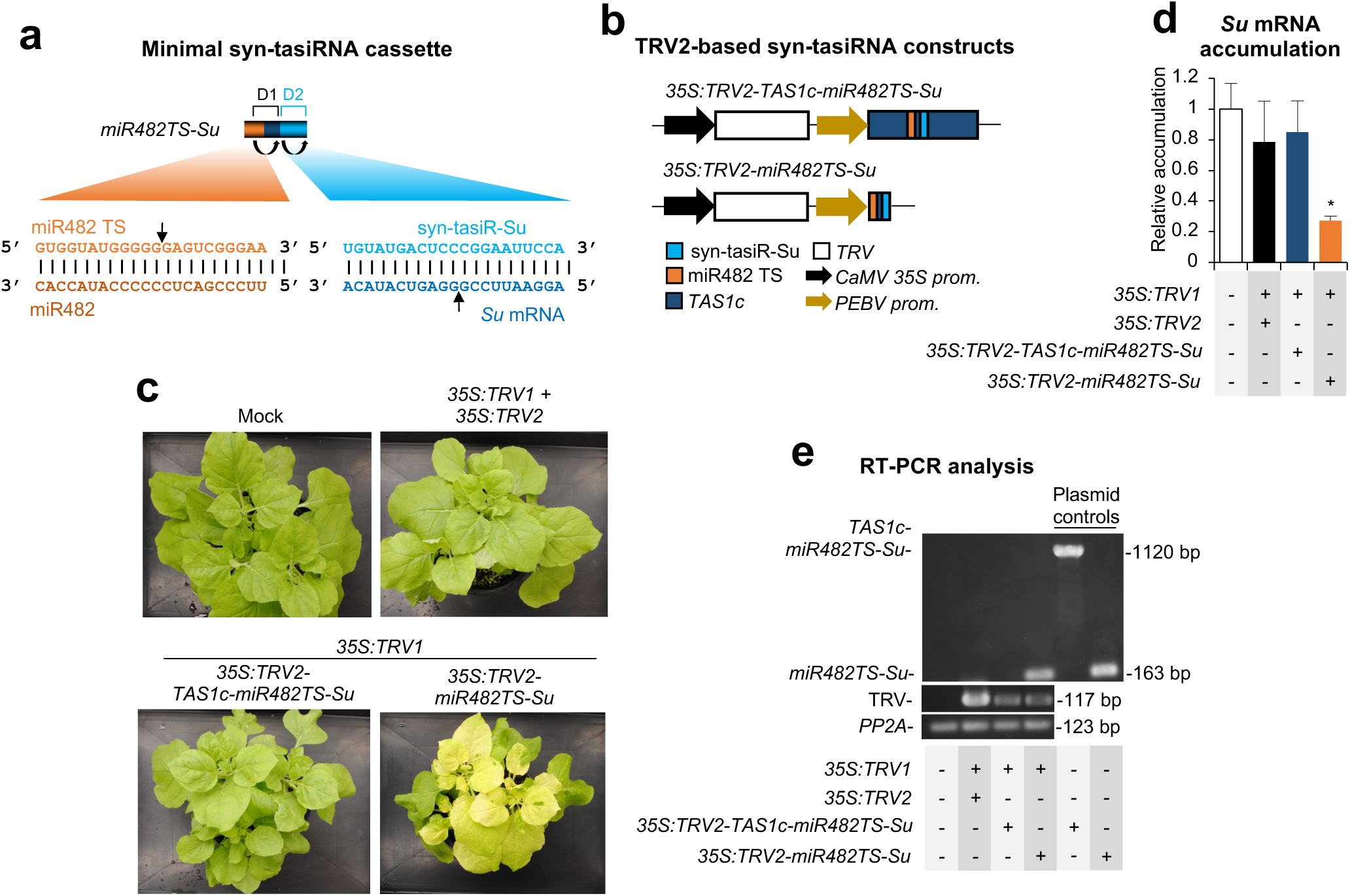
Functional analysis of tobacco rattle virus (TRV) constructs expressing syn-tasiR-Su from full-length *TAS1*c- or from minimal *miR482TS*-based syn-tasiRNA precursors in *Nicotiana benthamiana*. (**a**) Schematic representation of the anti-*Su* syn-tasiRNA cassette *miR482TS-Su*, engineered to express syn-tasiR-Su (light blue) from a minimal precursor containing the miR482 target site (TS) (orange) from *N. benthamiana* and a 11-nt spacer derived from *TAS1c* (dark blue). Other details are as described in Figure 1A. (**b**) Diagram of TRV-based constructs. *TAS1c*, miR482 target site (TS) and syn-tasiR-Su sequences are represented by dark blue, orange and light blue boxes, respectively. Other details are as in Figure 1B. (**c**) Photos at 14 days post-agroinoculation (dpa) of sets of three plants agroinoculated with the different constructs. (**d**) Target *Su* mRNA accumulation in RNA preparations from apical leaves collected at 7 dpa and analysed individually (mock = 1.0 in all comparisons). Bars with an asterisk indicate whether the mean values are significantly different from mock control samples (*P* < 0.05 in pairwise Student’s *t*-test comparison). (**e**) RT-PCR detection of TRV and syn-tasiRNA precursors in apical leaves at 7 dpa. Other details are as in Figure 1F.

Bleaching was first observed in certain areas of a few apical leaves at 8 dpa in plants agroinoculated with *35S:TRV2-miR482TS-Su*, and by 14-21 dpa it extended to most apical tissues (Figure 2C). In contrast, no bleaching was observed in plants agroinoculated with *35S:TRV2-TAS1c-miR482TS* or with control *35S:TRV2* at any time point (Figure 2C). RT-qPCR analysis confirmed that upper leaves from *35S:TRV2-miR482TS-Su*-expressing plants accumulated significantly lower levels of *Su* mRNA compared to controls (Figure 2D). RT-PCR analysis at 7 dpa of apical leaves detected the minimal *miR482TS-Su* precursor, whereas the full-length *TAS1c-miR482TS-Su* was not detected (Figure 2E). A TRV genomic fragment was detected in all TRV-treated plants, while *PP2a* was amplified in all samples (Figure 2E). Interestingly, Sanger sequencing of RT-PCR fragments from *TRV-miR482TS-Su*-infected plants revealed no mutations in the whole precursor insert.

Next, syn-tasiRNA biogenesis and processing from minimal *miR482TS*-based precursors in plants expressing *35S:TRV2-miR482TS* were analyzed in RNA preparations from upper leaves collected at 7 dpa. Northern blot analysis confirmed that syn-tasiR-Su accumulated predominantly as a single 21-nt band, whereas no signal was detected in mock-treated plants or control *GUS* plants (Figure 3A). High-throughput sequencing of sRNAs from RNA preparations of apical leaves confirmed that authentic syn-tasiR-Su was the predominant sRNA processed from the precursor, further validating its accurate processing *in planta* (Figure 3B).

**Figure 3.**
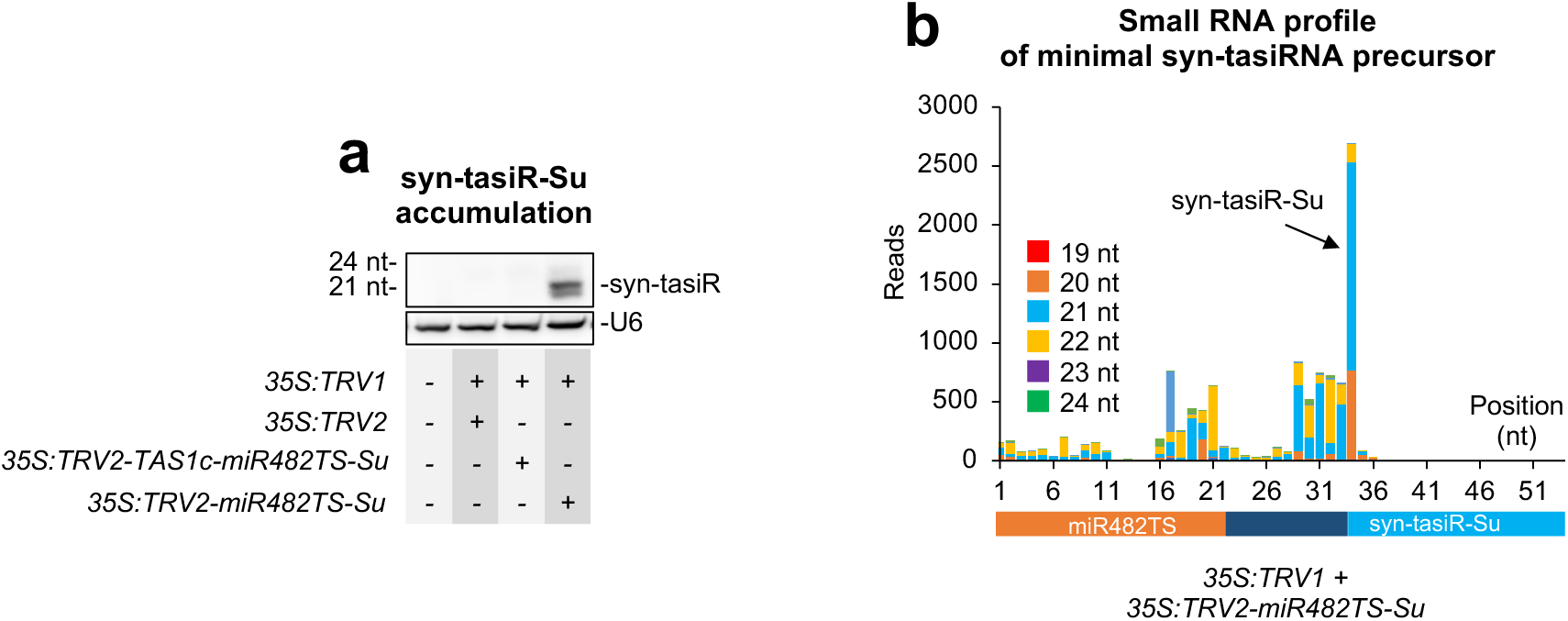
Accumulation and processing of syn-tasiR-Su expressed from *miR482TS*-based precursors in *Nicotiana benthamiana*. (**a**) Northern blot detection of syn-tasiR-Su in RNA preparations from apical leaves collected at 7 days post-agroinoculation (dpa). Other details are as described in Figure 1E. (**b**) sRNA profile of 19-24 nt [+] reads mapping to each of the 54 nucleotide positions within the *miR482aTS-Su* precursor from plants expressing *35S:TRV2-miR482TS-Su*. Orange, dark blue and light blue boxes represent nucleotides corresponding to *miR482TS*, the *TAS1c*-derived spacer and syn-tasiR-Su, respectively.

To rule out the possibility that *Su* silencing was due to siRNAs derived from the miR482TS-Su precursor originated during TRV replication, we co-agroinoculated plants with *35S:TRV1* and *35S:TRV2-miR173TS-Su*, which should not generate syn-tasiRNAs due to the absence of miR173 in *N. benthamiana* (Figure 4A). As controls, *35S:TRV1* was also independently agroinoculated with *35S:TRV2* and *35S:TRV2-miR482TS-Su* (Figure 4A). At 14 dpa, plants infiltrated with *35S:TRV2-miR482TS-Su* displayed widespread bleaching, as expected (Figure 4B). In contrast, plants expressing *35S:TRV2* or *35S:TRV2-miR173TS-Su* remained green, resembling mock-treated plants (Figure 4B). RT-qPCR analysis confirmed that *Su* mRNA levels were significantly reduced only in *35S:TRV2-miR482TS-Su*-expressing plants (Figure 4C). Finally, RT-PCR analysis in RNA samples extracted from apical leaves at 7 dpa confirmed the presence of the minimal precursors in *35S:TRV2-miR482TS-Su* and *35S:TRV2-miR173TS-Su* expressing plants (Figure 4D). TRV was present in all TRV-expressing plants, while *PP2A* was amplified in all samples, thus confirming that the lack of bleaching in *35S:TRV2-miR173a-Su*-expressing plants was not due to the deletion of the minimal precursor or to inefficient cDNA synthesis. Overall, these results support that TRV-based syn-tasiR-VIGS efficiently generates functional syn-tasiRNAs, and that *Su* silencing is specific and requires an endogenous 22-nt miRNA trigger to initiate syn-tasiRNA biogenesis.

**Figure 4.**
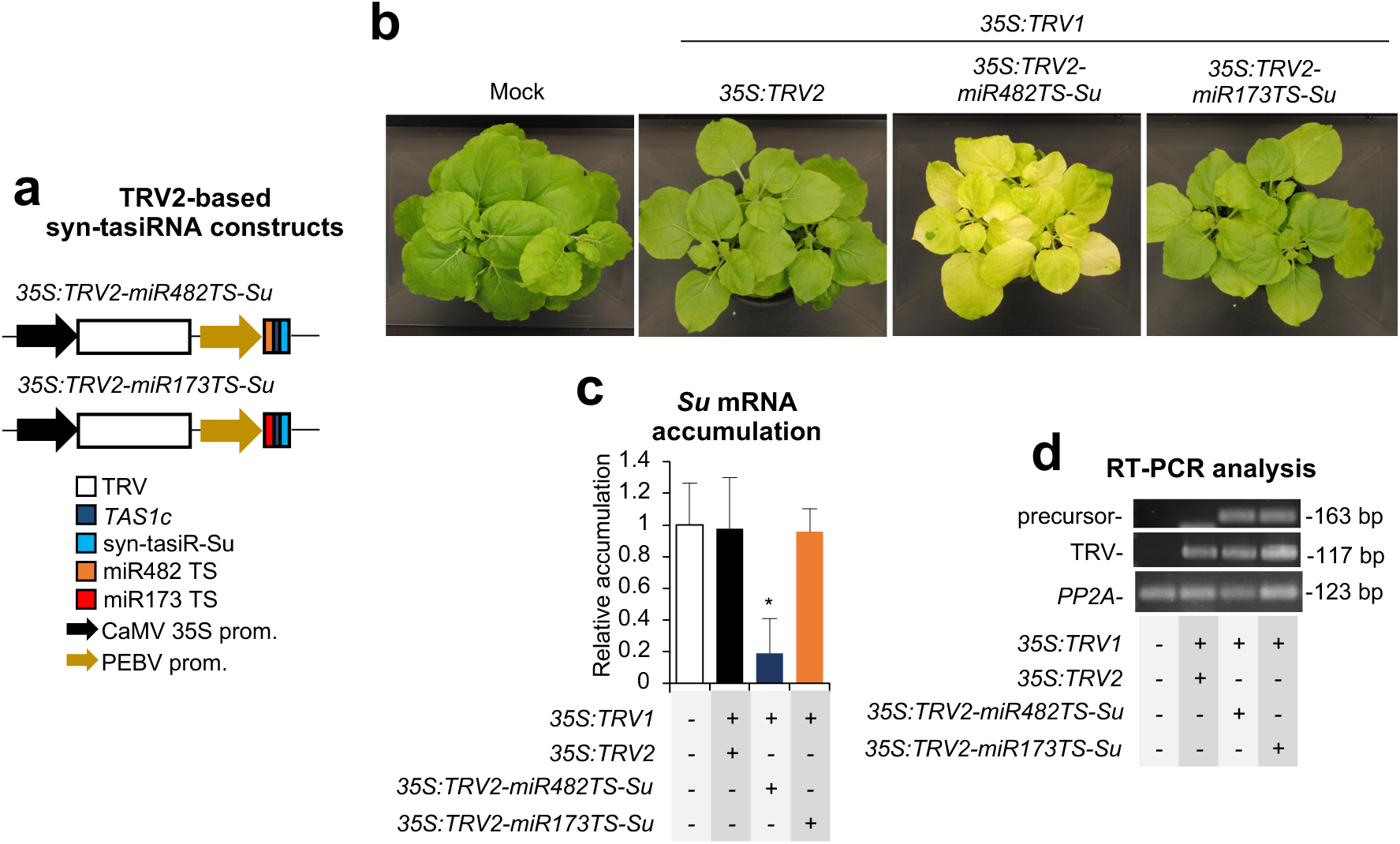
Functional analysis of tobacco rattle virus (TRV) constructs expressing syn-tasiR-Su minimal syn-tasiRNA precursors in *Nicotiana benthamiana*. (**a**) Diagram of TRV-based constructs. *TAS1c*, miR482 target site (TS), miR173 TS and syn-tasiR-Su sequences are represented by dark blue, orange, red and light blue boxes, respectively. Other details are as in Figure 1B. (**b**) Photos at 14 days post-agroinoculation (dpa) of sets of three plants agroinoculated with the different constructs. (**c**) Target *Su* mRNA accumulation in RNA preparations from apical leaves collected at 7 dpa and analysed individually (mock = 1.0 in all comparisons). Bar with an asterisk indicates whether the mean values are significantly different from mock control samples (*P* < 0.05 in pairwise Student’s *t*-test comparison). (**d**) RT-PCR detection of TRV and syn-tasiRNA precursors in apical leaves at 7 dpa. Other details are as in Figure 1F.

### Transgene-free, TRV-based syn-tasiR-VIGS for widespread gene silencing

Next, we aimed to establish TRV-based art-sRNA-VIGS as a non-transgenic, DNA-free gene silencing approach in *N. benthamiana*. The system involved two steps. First, several *N. benthamiana* plants were agroinoculated with *35S:TRV1* in combination with *35S:TRV2-shc-amiR-Su* or *35S:TRV2-miR482TS-Su* constructs, and after five days apical leaves were collected and crude extracts prepared (Figure 5A). Second, these TRV-containing crude extracts were sprayed onto young plants to assess transgene-free silencing of *Su*, as evidenced of bleaching phenotypes (Figure 5A). At 14 days post-spraying (dps), plants treated with TRV-shc-amiR-Su or TRV-miR482TS-Su crude extracts exhibited strong leaf bleaching phenotypes, while mock- and TRV-only-treated plants remained unaffected (Figure 5B). RT-PCR analysis confirmed the presence of TRV and minimal syn-tasiRNA precursors in apical leaves of TRV-shc-amiR-Su- and TRV-miR482TS-Su-treated plants at 14 dpa (Figure 5C). As expected, plants treated with TRV extracts accumulated TRV, whereas mock-treated plants lacked detectable TRV or minimal precursor signals (Figure 5C). *PP2A* was amplified in all samples (Figure 5C). Overall, these results indicate that TRV-based art-sRNA VIGS allows efficient, transgene-free gene silencing, offering a scalable and rapid tool for functional genomics.

**Figure 5.**
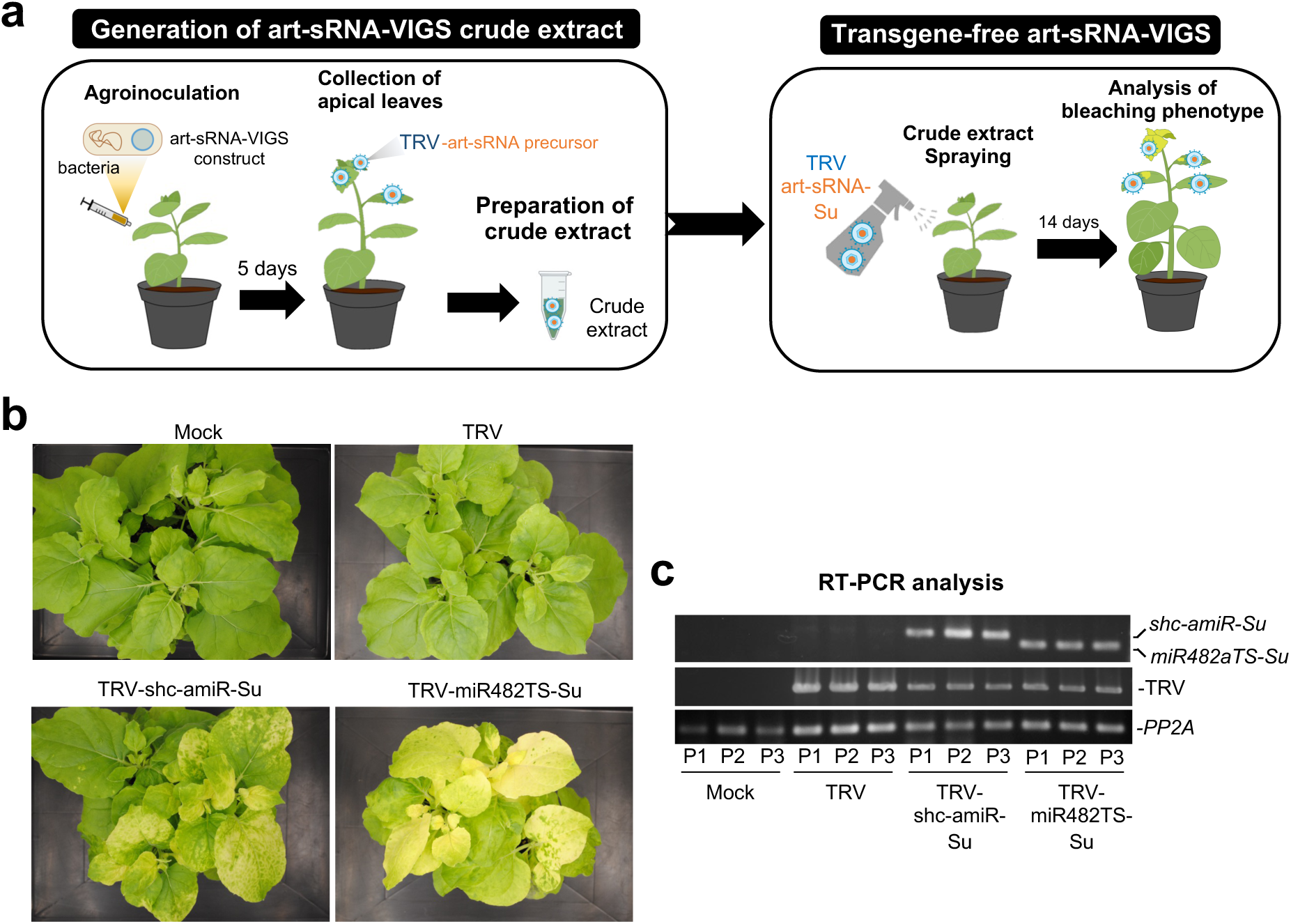
Non-transgenic, DNA-free widespread gene silencing in *Nicotiana benthamiana* through amiR-VIGS and syn-tasiR-VIGS using tobacco rattle virus (TRV). (**a**) Experimental set up to test gene silencing triggered by amiR-Su and syn-tasiR-Su expressed from TRV using minimal precursors. (**b**) Photos at 14 days post-spraying (dps) of sets of three plants sprayed with different crude extracts obtained from agroinoculated plants. (**c**) RT-PCR detection of TRV and minimal precursors in apical leaves at 14 dpa. Other details are as in Figure 1F.

### Plant resistance to a pathogenic virus by antiviral syn-tasiRNAs produced from TRV

To explore the potential of syn-tasiR-VIGS for antiviral resistance, we generated several TRV-based constructs expressing syn-tasiRNAs against TSWV. The *35S:TRV2-miR482TS-TSWV(x4)* construct included four validated anti-TSWV syn-tasiRNAs (syn-tasiR-TSWV-1, -2, -3, -4), previously shown to exhibit high antiviral activity (Carbonell et al. 2019), following miR482 TS (Figure 6A and B). Negative control constructs included *35S:TRV2-miR482TS-GUS(x4)*, expressing two syn-tasiRNAs (syn-tasiR-GUS-1 and syn-tasiR-GUS-2) targeting *GUS* (Carbonell, Lisón, et al., 2019) from miR482TS-based precursors, and *35S:TRV2-miR173TS-TSWV(x4)*, containing the miR173 TS but expected to be ineffective in triggering syn-tasiRNA biogenesis due to the absence of miR173 in *N. benthamiana* (Figure 6A and B).

**Figure 6.**
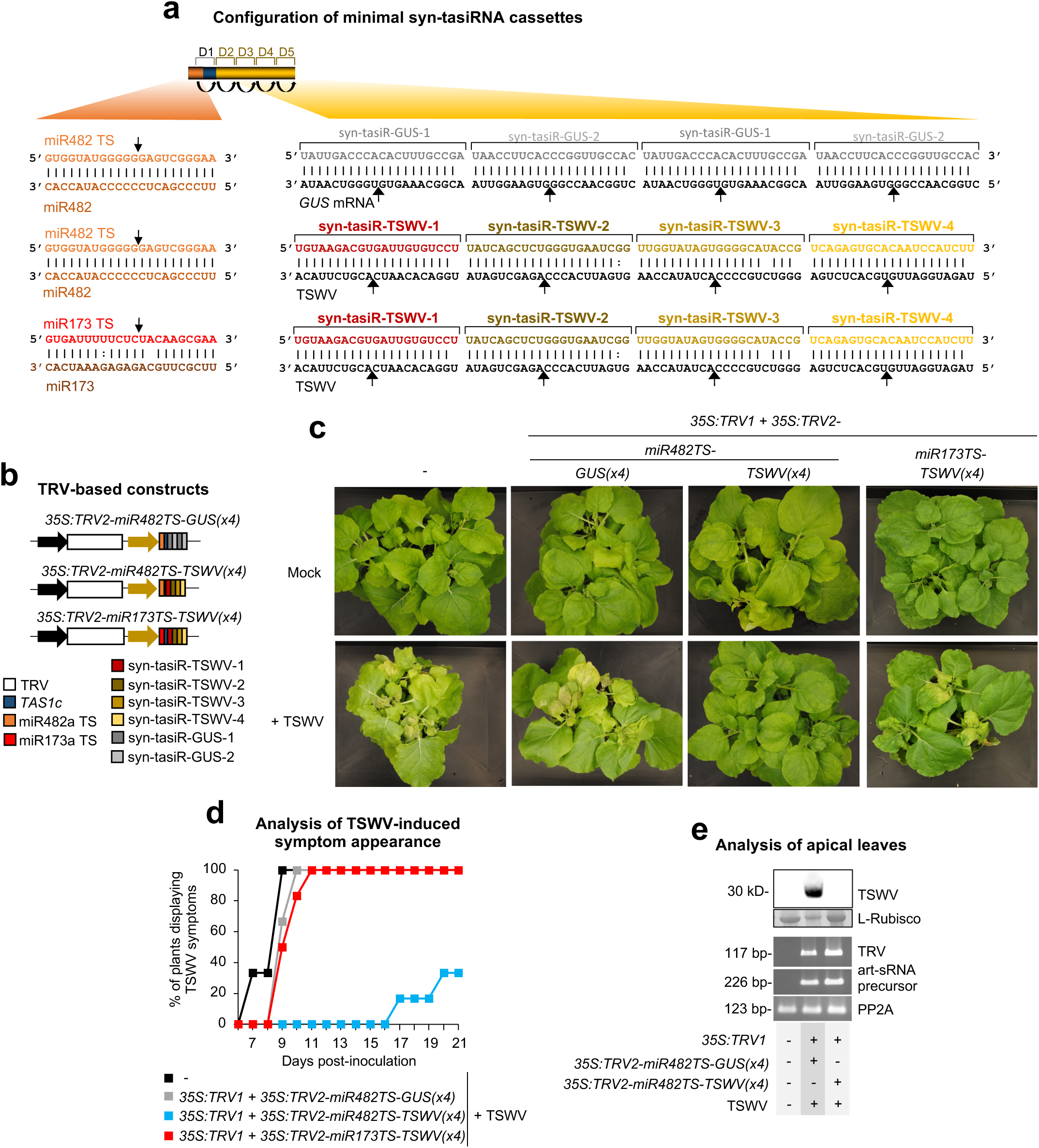
Functional analysis of tobacco rattle virus (TRV) constructs expressing syn-tasiRNAs against tomato spotted wilt virus (TSWV) in *Nicotiana benthamiana*. (**a**) Schematic representation of TRV-based constructs. Anti-TSWV art-sRNA sequences 1 (syn-tasiR-TSWV-1), 2 (syn-tasiR-TSWV-2), 3 (syn-tasiR-TSWV-3) and 4 (syn-tasiR-TSWV-4) are represented by red, dark brown, light brown and yellow boxes, respectively. Anti-GUS art-sRNA sequences 1 (syn-tasiR-GUS-1) and 2 (syn-tasiR-GUS-2) are represented by dark and light boxes, respectively. miR173 target site (TS) sequence is shown in a red box. Other details are as in Figure 2A. (**b**) Diagram of TRV-based constructs.. (**c**) Photos at 21 days post-inoculation (dpi) of sets of three plants agroinoculated with the different constructs and inoculated or not (mock) with TSWV. (**d**) Two-dimensional line graph showing, for each of the six-plant sets listed, the percentage of symptomatic plants per day during 21 days. (**e**) Analysis of apical leaves collected at 14 dpi and pooled from six independent plants. Top, Western blot detection of TSWV in protein preparations. The membrane stained with Ponceau red showing the large subunit of Rubisco (ribulose1,5biphosphate carboxylase/oxygenase) is included as loading control. Bottom, RT-PCR detection of TRV and syn-tasiRNA precursors. Other details are as in Figure 1F.

To assess the antiviral activity of TRV-based constructs, each construct was agroinoculated into one leaf of six independent *N. benthamiana* plants. After five days, these plants were further inoculated with TSWV, and symptom progression was monitored over 28 days. By 14 days post-inoculation (dpi), all plants expressing anti-TSWV syn-tasiRNAs from *miR482TS* precursor remained asymptomatic, whereas all control plants, including those expressing syn-tasiR-GUS and *miR173TS* precursors, developed severe TSWV symptoms (Figure 6C and 6D). At this same time point, Western blot analysis of apical leaves showed that none of the plants expressing *35S:TRV2-miR482TS-TSWV(x4)* accumulated TSWV, while control plants expressing *35S:TRV2-miR482TS-GUS(x4*) exhibited high TSWV accumulation (Figure 6E). RT-PCR analysis confirmed the presence of a 226-bp fragment corresponding to the minimal precursors and a 117-bp fragment from the TRV genome in all TRV-treated samples, whereas these fragments were absent in mock-inoculated and non-agroinfiltrated plants (Figure 6E). By 28 dpi, four out of six plants expressing anti-TSWV syn-tasiRNAs remained completely symptom-free (Figure 6D).

sRNA blot analysis at 7 dpa in *35S:TRV2-miR482TS-TSWV(x4)* non-inoculated plants confirmed the presence of high levels of 21-nt anti-TSWV syn-tasiRNAs, whereas no corresponding signals were detected in control samples (Figure 7A). The accuracy of *miR482TS-TSWV(x4)* precursor processing and the production of authentic anti-TSWV syn-tasiRNAs were analyzed by high-throughput sequencing of sRNA libraries from *N. benthamiana* plants agroinoculated with *35S:TRV2-miR482-TSWV(x4)* (Figure 7B). All four authentic syn-tasiRNA sequences were detected *in vivo*, although at varying levels. Syn-tasiR-TSWV-1 and syn-tasiR-TSWV-2 syn-tasiRNAs were detected as predominant 21-nt sequences when plotting all 19-24 (+) sRNAs mapping to the precursor, while syn-tasiR-TSWV-3 and syn-tasiR-TSWV were detected at lower levels (Figure 7B). Additionally, phasing analysis showed that 46% of 21-nt [+] reads aligning to the first register (Figure 7B), confirming precise processing of the precursor. Taken together, these results indicate that TRV-based syn-tasiR-VIGS effectively triggers the production of multiple syn-tasiRNAs *in planta* and further highlight the potential of RNA viral vectors like TRV to deliver syn-tasiRNAs and confer complete antiviral immunity in plants.

**Figure 7.**
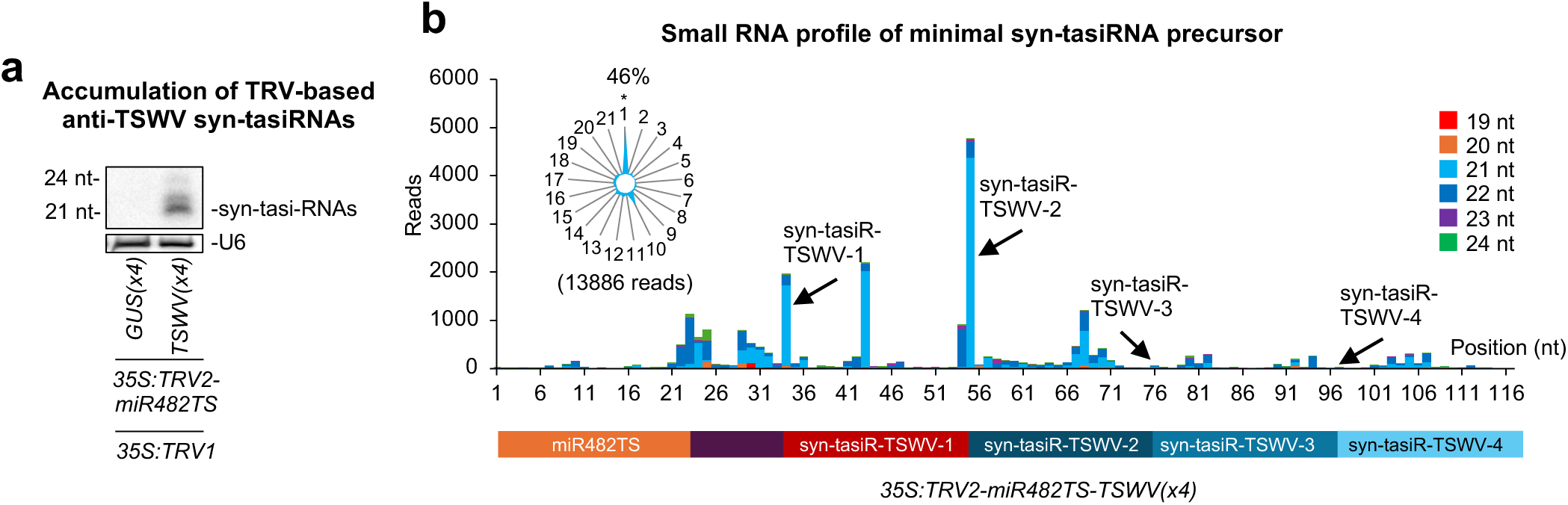
Accumulation and processing of anti-TSWV syn-tasiRNAs expressed from *miR482TS*-based precursors in *Nicotiana benthamiana*. (**a**) Northern blot detection of anti-TSWV art-sRNAs in RNA preparations from apical leaves collected at 7 dpa and pooled from three independent mock-inoculated plants. A cocktail of probes to simultaneously detect syn-tasiR-TSWV-1, syn-tasiR-TSWV-2, syn-tasiR-TSWV-3 and syn-tasiR-TSWV-4 was used. (**b**) syn-tasiRNA processing from TRV-miR482TS-TSWV(x4). Top: sRNA profile of 19-24 nt [+] reads mapping to each of the 117 nucleotide positions in the *miR482TS-TSWV(x4)* precursor from samples expressing *35S:TRV2-NbmiR482TS-TSWV(x4)*. Radar plot shows the proportion of 21-nt reads corresponding to each of the 21 registers from the minimal syn-tasiRNA precursor, with position 1 designated as immediately after miR482-guided cleavage site. Other details are as in Figure 6A.

## DISCUSSION

Here, we show that a TRV-based viral vector can efficiently produce amiRNAs and syn-tasiRNAs from minimal precursors to induce widespread gene silencing and antiviral resistance in *N. benthamiana*. These results highlight the unique advantages of minimal precursors and extend the application of art-sRNA-VIGS beyond previously established viral vector systems (Tang et al. 2010; Ju et al. 2017; Kuo and Falk 2022; Cisneros et al. 2023; Cisneros et al. 2025).

TRV has been an ideal viral vector for gene silencing in *N. benthamiana* and other plant species since the early 2000’s (Ratcliff et al. 2001; Lu et al. 2003; Burch-Smith et al. 2004). Unlike other RNA viruses, TRV induces minimal or no symptoms in *N. benthamiana*, preventing confounding effects of viral pathogenicity on plant phenotypes (Ratcliff et al. 2001). Moreover, its efficient systemic movement allows widespread gene silencing, while its bipartite genome permits flexible engineering without compromising viral replication or silencing efficiency. These features make TRV particularly suited for delivering minimal amiRNA and syn-tasiRNA precursors to plants in an effective and symptom-free manner. Additionally, the recently developed JoinTRV expression system, based on mini T-DNA vectors with compatible origins (Aragonés et al. 2022), facilitates the simple, one-step cloning of short double-stranded DNA inserts including minimal precursor sequences into *pLX-TRV2* (Text S1), streamlining the generation of art-sRNA-VIGS constructs in a time- and cost-effective manner.

A key feature of TRV-based art-sRNA-VIGS is the use of minimal precursors, which enhance stability during viral replication while reducing the accumulation of mutations. Here, both *pri*- and *TAS1c*-based full-length precursors were rapidly deleted from TRV a few days after infection, whereas minimal precursors were stably maintained and yielded high levels of functional amiRNAs and syn-tasiRNAs. The deletion of full-length precursors underscores the limited cargo capacity of viral vectors for VIGS (Rössner et al. 2022) and highlights the need for minimal designs to ensure precursor retention. Our results are consistent with previous studies using PVX-based art-sRNA VIGS, which also showed that only minimal precursors efficiently generated functional art-sRNAs *in planta* (Cisneros et al. 2023; Cisneros and Carbonell 2025). Remarkably, TRV-based art-sRNAs were accurately processed from minimal precursors and accumulated to high levels, as confirmed by high-throughput sRNA sequencing and northern blot. Since TRV replicates in the cytoplasm, and DCL4 is the main DCL functioning in antiviral defense against RNA viruses –producing 21-nt sRNAs from diced viral RNAs (Deleris et al. 2006; Bouche et al. 2006), it is tempting to speculate that DCL4 is responsible for processing amiRNA precursors present in TRV. Indeed, genetic analyses using DCL-RNAi knockdown plants supported that DCL4 is involved in amiRNA processing from PVX-based viral vectors (Cisneros et al. 2023). Curiously, DCL4, in addition to processing long dsRNA precursors, can access flexibly structured single-stranded RNA substrates such as pre-miRNA-like RNAs, as shown in the biogenesis of several sRNAs produced from cucumber mosaic virus satellite RNA (Du et al. 2007). Conversely, the involvement of DCL4 in syn-tasiRNA biogenesis from TRV is expected following RDR6/SGS3-mediated dsRNA synthesis after NbmiR482a cleavage.

Another key feature of art-sRNA VIGS is its high specificity. Unlike classic dsRNA-based VIGS, which generates heterogenous populations of siRNAs with potential off-target effects (Burch-Smith et al. 2004), art-sRNAs are computationally designed to ensure precise targeting while minimizing unintended molecular interactions (Ossowski et al. 2008; Fahlgren et al. 2016). Our high-throughput sequencing data confirm that TRV-expressed minimal precursors are accurately processed in *N. benthamiana*, yielding authentic art-sRNAs that accumulate at high levels relative to other sRNAs derived from the precursor (Figure 3B and 7B). Moreover, no phased secondary 21-nt sRNAs derived from *Su* were detected in plants expressing syn-tasiR-Su (Data S2, Figure S1), reinforcing the specificity of the TRV-based art-sRNA-VIGS approach. Importantly, the absence of bleaching and resistance in plants expressing *35S:TRV2-miR173TS-Su* and *35S:TRV2-miR173aTS-TSWV(x4)*, respectively, supports the conclusion that gene silencing results from syn-tasiRNA activity rather than from potential siRNAs generated during TRV replication.

Another important finding is the successful adaptation of TRV-based art-sRNA VIGS for transgene-free gene silencing through crude extract spraying, as shown recently for PVX-based amiR-VIGS and syn-tasiR-VIGS (Cisneros et al. 2023; Cisneros and Carbonell 2025). This strategy eliminates the need for genetic transformation, enabling rapid and scalable functional studies in plants. The ability to deliver TRV-based art-sRNA VIGS in a non-transgenic manner broadens its potential applications in both research and agriculture. Future research should advance art-sRNA-VIGS by exploring viral vectors with broader or species-specific host ranges, particularly for application in crops. However, several challenges remain in implementing VIGS for crop improvement: i) variable efficiency across species, tissues, and environments, ii) unintended phenotypic changes due to off-target effects, iii) potential growth retardation and yield reduction, iv) vector instability during prolonged infections, and v) biosafety concerns over the environmental release of genetically modified viral vectors. Addressing these limitations will require the development of alternative approaches, such as topical art-sRNA delivery or CRISPR/Cas-editing of endogenous sRNA loci.

In conclusion, our study establishes TRV-based art-sRNA VIGS as a functional and versatile RNAi platform in *N. benthamiana* based on TRV, offering a new tool for highly specific gene silencing in both transient and transgene-free applications. Further adaptation of TRV-based art-sRNA-VIGS to additional viral vector systems and plant species is necessary to maximize its potential. While TRV offers advantages such as systemic spread and minimal symptomatology, other RNA viral vectors with different host ranges and characteristics may be better suited for specific applications. Future research should explore art-sRNA-VIGS within alternative viral platforms, particularly those compatible with economically important crops. Additionally, optimizing minimal precursor designs to optimize processing efficiency and target specificity will further improve the applicability of this approach.

## Supplementary Information

The online version contains supplementary material available at https://doi. org/xx.xxx/xxxxxx-xxx-xxx-x.

## Authors’ contributions

MJ-M did most of the experimental work with the help of AA, AP, IO-M and AEC. MJ-M, AEC and AC analyzed the data. A.C. conceived the research, supervised the project and wrote the manuscript with input from the rest of authors.

## Availability of data and materials

All data generated or analyzed during this study are included in this published article and its supplementary information files. High-throughput sequencing data can be found in the Sequence Read Archive (SRA) database under accession number PRJNA1241532.

## ACKNOWLEDGEMENTS

We are grateful to José-Antonio Daròs (IBMCP) for providing the *pLB-TRV1* and *pLB-TRV2* plasmids, Javier Forment (IBMCP) for his support with the analysis of sRNA sequencing data, Sara Toledano for excellent technical assistance, and the greenhouse staff at IBMCP for their help in plant growth and maintenance.

