## Supplementary material for "Gene Silencing in Plants by Artificial Small RNAs Derived from Minimal Precursors and Expressed via Tobacco Rattle Virus": Figure S1

### Slide 1
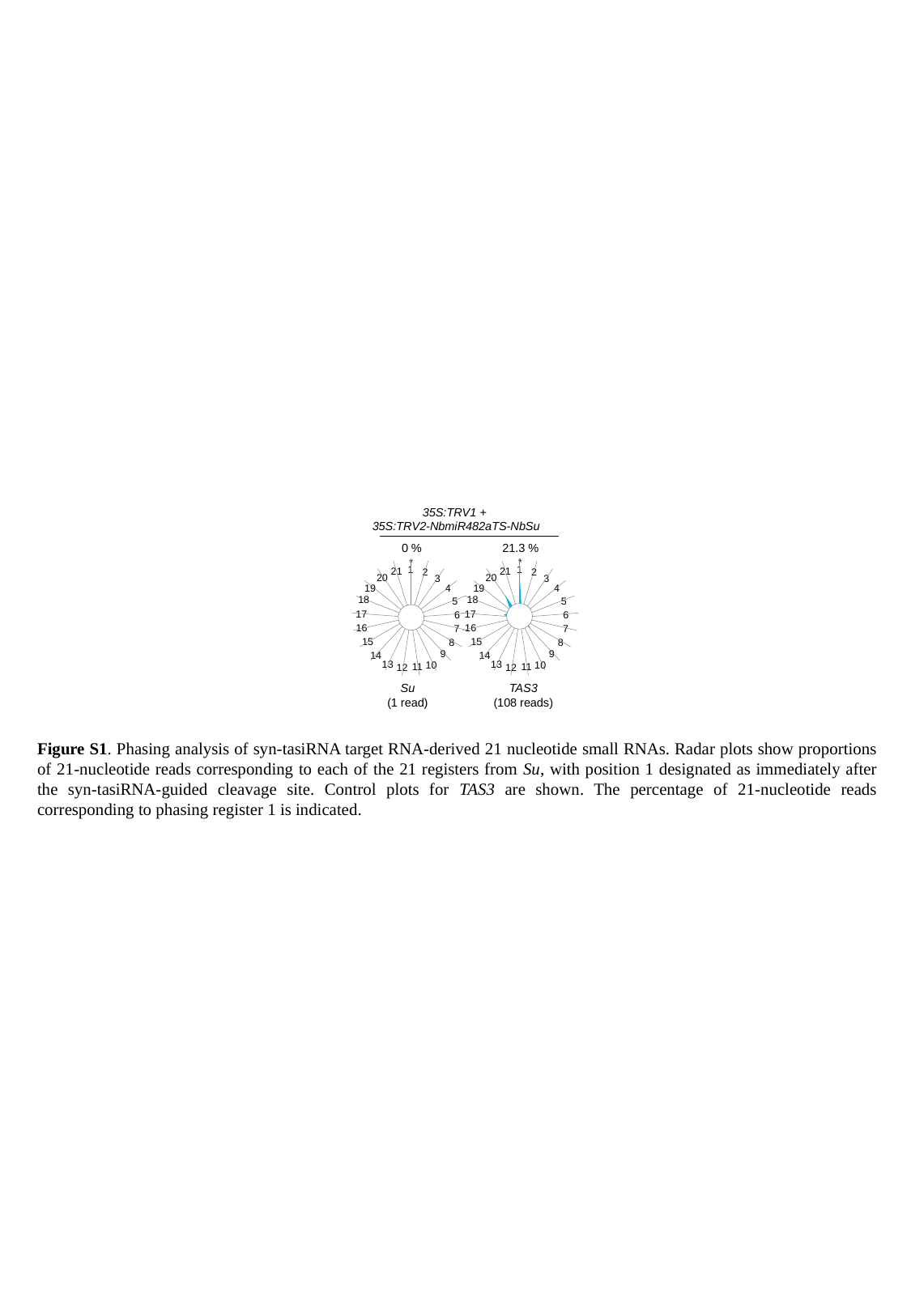

35S:TRV1 +
35S:TRV2-NbmiR482aTS-NbSu
#### Chart
| Category | |
|---|---|0 %
*
#### Chart
| Category | |
|---|---|1
21
2
20
3
4
19
18
5
#### Chart
| Category | |
|---|---|17
6
16
7
15
8
9
14
13
10
11
12
Su
(1 read)
21.3 %
*
#### Chart
| Category | |
|---|---|1
21
2
20
3
4
19
18
5
#### Chart
| Category | |
|---|---|17
6
16
7
15
8
9
14
13
10
11
12
TAS3
(108 reads)
Figure S1. Phasing analysis of syn-tasiRNA target RNA-derived 21 nucleotide small RNAs. Radar plots show proportions of 21-nucleotide reads corresponding to each of the 21 registers from Su, with position 1 designated as immediately after the syn-tasiRNA-guided cleavage site. Control plots for TAS3 are shown. The percentage of 21-nucleotide reads corresponding to phasing register 1 is indicated.
