## Supplementary material for "Gene Silencing in Plants by Artificial Small RNAs Derived from Minimal Precursors and Expressed via Tobacco Rattle Virus": Table S1

**Table S1.** Name, sequence and use of DNA oligonucleotides used in this study.

| **Name** | **Sequence** | **Type*** | **Construct/Aim** |
| --- | --- | --- | --- |
| AC-50 | CCGATTCACCCAGAGCTGATA | ssDNA | Probe to detect syn-tasiR-TSWV-2 |
| AC-51 | CGGTATGCCCCACTATACCAA | ssDNA | Probe to detect syn-tasiR-TSWV-3 |
| AC-52 | AAGATGGATTGTGCACTCTGA | ssDNA | Probe to detect syn-tasiR-TSWV-4 |
| AC-55 | AGGGGCCATGCTAATCTTCTC | ssDNA | Probe for U6 detection |
| AC-355 | GACCCTGATGTTGATGTTCGCT | ssDNA | qPCR amplification of *Su* mRNA |
| AC-356 | GAGGGATTTGAAGAGAGATTTC | ssDNA |  |
| AC-365 | GACCCTGATGTTGATGTTCGCT | ssDNA | PCR&qPCR amplification of *PP2A* mRNA |
| AC-366 | GAGGGATTTGAAGAGAGATTTC | ssDNA |  |
| AC-416 | A+GGA+CAC+AAT+CAC+GTC+TTA+CA | ssLNA | Probe to detect syn-tasiR-TSWV-1 |
| AC-417 | G+CGG+GAA+GTC+CAC+CAC+GGT+TA | ssLNA | Probe for amiR-Su/syn-tasiR-Su detection |
| AC-518 | cacttacccgagttaacgccAAACCTAAACCTAAACGG | ssDNA | *35S:TRV2-TAS1c-miR482TS-Su* |
| AC-519 | gtttaatgtcttcgggacatATTTCACTTTACGATGTGG | ssDNA |  |
| AC-523 | TCGGTTTGCTGACCTACTGG | ssDNA | art-sRNA precursor detection |
| AC-524 | AACCTAAAACTTCAGACACGG | ssDNA |  |
| AC-615 | cacttacccgagttaacgccTATAGGGGGGAAAAAAAGGTAG | ssDNA | *35S:TRV2-pri-amiR-Su* |
| AC-616 | gtttaatgtcttcgggacatGAGACTAAAGATGAGATCTAATCTG | ssDNA |  |
| AC-617 | cacttacccgagttaacgccAGTAGAGAAGAATCTGTA | ssDNA | *35S:TRV2-shc-amiR-Su* |
| AC-618 | gtttaatgtcttcgggacatAGTAAGAAGAGCCAA | ssDNA |  |
| AC-660 | ATGGGAGATATGTACGATGAAT | ssDNA | TRV diagnostic |
| AC-661 | GGGATTAGGACGTATCGGACC | ssDNA |  |
| AC-667 | cacttacccgagttaacgccGTGGTATGGGGGGAGTCGGGAATAGACCATTTATGTATGACTCCCGGAATTCCAatgtcccgaagacattaaac | dsDNA | *35S:TRV2-miR482TS-Su* |
| AC-984 | cacttacccgagttaacgccGTGGTATGGGGGGAGTCGGGAATAGACCATTTATATTGACCCACACTTTGCCGATAACCTTCACCCGGTTGCCACTATTGACCCACACTTTGCCGATAACCTTCACCCGGTTGCCACatgtcccgaagacattaaac | dsDNA | *35S:TRV2-miR482TS-GUS(x4)* |
| AC-985 | cacttacccgagttaacgccGTGGTATGGGGGGAGTCGGGAA | ssDNA | *35S-TRV2-miR482TS-TSWV(x4)* |
| AC-986 | gtttaatgtcttcgggacatAAGATGGATTGTGCACTCTGA | ssDNA |  |
| AC-1222 | cacttacccgagttaacgccGTGATTTTTCTCTACAAGCGAA | ssDNA | *35S:TRV2-miR173TS-Su* |
| AC-1223 | gtttaatgtcttcgggacatTGGAATTCCGGGAGTCATACATAAA | ssDNA |  |

*ssDNA: single-stranded DNA; dsDNA: double-stranded DNA; LNA: locked nucleic acid.
