## Supplementary material for "Gene Silencing in Plants by Artificial Small RNAs Derived from Minimal Precursors and Expressed via Tobacco Rattle Virus": Text S1

**Text S1.** Protocol to generate TRV-based art-sRNA constructs.

**1. Preparation of the dsDNA art-sRNA insert**

**1.1. amiRNA insert**

Design and order a dsDNA (eg. ultramer duplex in IDT) including the sequence of your amiRNA included in the *MIR390*-based ("*shc*") minimal precursor, as follows:

cacttacccgagttaacgccAGTAGAGAAGAATCTGTA**X_1_X_2_X_3_X_4_X_5_X_6_X_7_X_8_X_9_X_10_X_11_X_12_X_13_X_14_X_15_X_16_X_17_X_18_X_19_X_20_X_21_AGTAGAGAAGAATCTGTAX_1_X_2_X_3_X_4_X_5_X_6_X_7_X_8_X_9_X_10_X_11_X_12_X_13_X_14_X_15_X_16_X_17_X_18_X_19_X_20_X_21_**CGAAATCAAACT**X_1_X_2_X_1_X_2_X_3_X_4_X_5_X_6_X_7_X_8_X_9_X_10_X_11_X_12_X_13_X_14_X_15_X_16_X_17_X_18_X_19_CATTGGCTCTTCTTACT**atgtcccgaagacattaaac

Where:

-**X** is a DNA base of the amiRNA sequence, and the subscript number is the base position in the amiRNA 21-mer

-**X** is a DNA base of the amiRNA* sequence, and the subscript number is the base position in the amiRNA* 21-mer

-**X** is a DNA base of the basal stem (BS) region of the *shc* precursor

-X is a DNA base of the *shc* precursor included in the oligonucleotides required to clone the amiRNA insert in B/c vectors

-x is a DNA base of the TRV sequence, required for Gibson-based assembly

-**X** is a DNA base of the *shc* precursor included in the oligonucleotides required to clone the amiRNA insert in B/c vectors

-**X** is a DNA base of the *shc* precursor that may be modified to preserve the authentic *shc* duplex structure

In the sequence above:

-Insert the amiRNA sequence where you see **X_1_X_2_X_3_X_4_X_5_X_6_X_7_X_8_X_9_X_10_X_11_X_12_X_13_X_14_X_15_X_16_X_17_X_18_X_19_X_20_X_21_**

-Insert the amiRNA* sequence that has to verify the following base-pairing:

**X_1_ X_2_ X_3_ X_4_ X_5_ X_6_ X_7_ X_8_ X_9_ X_10_X_11_X_12_X_13_X_14_X_15_X_16_X_17_X_18_ X_19_ X_20_X_21_**	**|**	**|	|	|**	**|	|	|**	**|	|	|**	**|	|**	**|	|	|**	**|	|	|**	**|	|**

**X_19_X_18_X_17_X_16_X_15_X_14_X_13_X_12_X_11_X_10_X_9_ X_8_ X_7_ X_6_ X_5_ X_4_ X_3_ X_2_ X_1_ X_2_ X_1_**

Note that:

-In general, **X_1_=T** for amiRNA association with AGO1. In this case, **X_19_=A**

-Bases **X_11_** and **X_9_** DO NOT base-pair to preserve the central bulge of the authentic *AtMIR390a* duplex. The following base-pair rule applies:

-If **X_11_**=G, then **X_9_**=A

-If **X_11_**=C, then **X_9_**=T

-If **X_11_**=A, then **X_9_**=G

-If **X_11_**=U, then **X_9_**=C

Fragment #1 (amiRNA precursor) is ready.

**1.2. syn-tasiRNA insert**

Design and order a dsDNA (eg. ultramer duplex in IDT) including the sequences of your syn-tasiRNA(s) (2 in the following example) following the 22-nt miRNA target site of interest, as follows:

agaggtcagcaccagctagc**X_1_X_2_X_3_X_4_X_5_X_6_X_7_X_8_X_9_X_10_X_11_X_12_X_13_X_14_X_15_X_16_X_17_X_18_X_19_X_20_X_21_X_22_**TAGACCATTTA**X_1_X_2_X_3_X_4_X_5_X_6_X_7_X_8_X_9_X_10_X_11_X_12_X_13_X_14_X_15_X_16_X_17_X_18_X_19_X_20_X_21_X_1_X_2_X_3_X_4_X_5_X_6_X_7_X_8_X_9_X_10_X_11_X_12_X_13_X_14_X_15_X_16_X_17_X_18_X_19_X_20_X_21_**agggtttgttaagtttccct

Where:

-**X** is a DNA base of the 22-nt miRNA target site sequence, and the subscript number is the base position

-**X** is a DNA base of the syn-tasiRNA-1 sequence, and the subscript number is the base position in the syn-tasiRNA* 21-mer

-**X** is a DNA base of the syn-tasiRNA-2 sequence, and the subscript number is the base position in the syn-tasiRNA 21-mer

-x is a DNA base of the TRV sequence, required for Gibson-based assembly

-X is a DNA base of the *AtTAS1c* sequence

Note that:

-In general, **X_1_=T** and **X_1_=T** for amiRNA association with AGO1.

Fragment #1 (syn-tasiRNA precursor) is ready.

**2. Preparation of the vector**

-Digest *pLB-PVX* with *Mlu*I.

-Gel purify the 9921 bp band corresponding to linearized plasmid.

-Quantify 1 ul in Nanodrop.

Fragment #2 (backbone vector) is ready.

**3. Assembly**

-Assemble the Gibbson reaction as described below:

Fragment 1 (dsDNA insert)^a^

Fragment 2 (vector)^b,c,d^

GeneArt Gibson Assembly HiFI Master Mix	5 μL

dH_2_O	to 10 μL

Total volume	10 μL

^a^The optimal amount of vector is between 50-100 ng

^b^Insert/vector molar excess is between 2-3.

^c^Total DNA amount is between 0.02-0.5 pmol

^d^Mass to moles conversions can be calculated here:

<http://nebiocalculator.neb.com/#!/ssdnaamt>

-Incubate reactions at 50ºC for 1h.

-Clean up reactions with a column (e.g. Zymo Research)

-Transform 1-4 μL in *E. coli* DH5α

-Plate in L-Kan plates and incubate 16h at 37ºC

**4. Clone verification**

-Pick several colonies and grow in liquid LB-Kan 16h at 37ºC, and purify plasmids.

-Digest candidate clones with *BpiI*

Good clones:

1 amiRNA: 2296 + 1064 + **1011 +** 1064 + 728 + 423 + 378 +3 bp

2 syn-tasiRNAs: 2296 **+** 1064 + **997** + 728 + 423 + 378 +3 bp

Bad clones (empty *pLX-TRV2*): 2296 + **1351 +** 1064 + 728 + 423 + 378 +3 bp

-Confirm insert sequence by Sanger sequencing with forward and reverse oligos AC-523 (TCGGTTTGCTGACCTACTGG) and/or AC-524 (AACCTAAAACTTCAGACACGG), respectively.
