## Supplementary material for "Gene Silencing in Plants by Artificial Small RNAs Derived from Minimal Precursors and Expressed via Tobacco Rattle Virus": Text S2

**Text S2**. DNA sequence in FASTA format of all precursors used to express art-sRNAs in plants.

1. amiRNA precursors

***>pri-amiR-Su***

TATAGGGGGGAAAAAAAGGTAGTCATCAGATATATATTTTGGTAAGAAAATATAGAAATGAATAATTTCACGTTTAACGAAGAGGAGATGACGTGTGTTCCTTCGAACCCGAGTTTTGTTCGTCTATAAATAGCACCTTCTCTTCTCCTTCTTCCTCACTTCCATCTTTTTAGCTTCACTATCTCTCTATAATCGGTTTTATCTTTCTCTAAGTCACAACCCAAAAAAACAAAGTAGAGAAGAATCTGTATGTATGACTCCCGGAATTCCAATGATGATCACATTCGTTATCTATTTTTTTGGAATTCCGTGAGTCATACACATTGGCTCTTCTTACTACAATGAAAAAGGCCGAGGCAAAACGCCTAAAATCACTTGAGAATCAATTCTTTTTACTGTCCATTTAAGCTATCTTTTATAAACGTGTCTTATTTTCTATCTCTTTTGTTTAAACTAAGAAACTATAGTATTTTGTCTAAAACAAAACATGAAAGAACAGATTAGATCTCATCTTTAGTCTC

*AtMIR390a*

amiR-Su

amiR-Su*

***>shc-amiR-Su***

AGTAGAGAAGAATCTGTATGTATGACTCCCGGAATTCCACGAAATCAAACTTGGAATTCCGTGAGTCATACACATTGGCTCTTCTTACT

*AtMIR390a*

*OsMIR390*

amiR-Su

amiR-Su*

2. syn-tasiRNA precursors

**>*TAS1c-Su***

AAACCTAAACCTAAACGGCTAAGCCCGACGTCAAATACCAAAAAGAGAAAAACAAGAGCGCCGTCAAGCTCTGCAAATACGATCTGTAAGTCCATCTTAACACAAAAGTGAGATGGGTTCTTAGATCATGTTCCGCCGTTAGATCGAGTCATGGTCTTGTCTCATAGAAAGGTACTTTCGTTTACTTCTTTTGAGTATCGAGTAGAGCGTCGTCTATAGTTAGTTTGAGATTGCGTTTGTCAGAAGTTAGGTTCAATGTCCCGGTCCAATTTTCACCAGCCATGTGTCAGTTTCGTTCCTTCCCGTCCTCTTCTTTGATTTCGTTGGGTTACGGATGTTTTCGAGATGAAACAGCATTGTTTTGTTGTGATTTTTCTCTACAAGCGAATAGACCATTTATGTATGACTCCCGGAATTCCATCGGTGGATCTTAGAAAATTATTCTAAGTCCAACATAGCGTATTCTAAGTTCAACATATCGACGAACTAGAAAAGACATTGGACATATTCCAGGATATGCAAAAGAAAACAATGAATATTGTTTTGAATGTGTTCAAGTAAATGAGATTTTCAAGTCGTCTAAAGAACAGTTGCTAATACAGTTACTTATTTCAATAAATAATTGGTTCTAATAATACAAAACATATTCGAGGATATGCAGAAAAAAAGATGTTTGTTATTTTGAAAAGCTTGAGTAGTTTCTCTCCGAGGTGTAGCGAAGAAGCATCATCTACTTTGTAATGTAATTTTCTTTATGTTTTCACTTTGTAATTTTATTTGTGTTAATGTACCATGGCCGATATCGGTTTTATTGAAAGAAAATTTATGTTACTTCTGTTTTGGCTTTGCAATCAGTTATGCTAGTTTTCTTATACCCTTTCGTAAGCTTCCTAAGGAATCGTTCATTGATTTCCACTGCTTCATTGTATATTAAAACTTTACAACTGTATCGACCATCATATAATTCTGGGTCAAGAGATGAAAATAGAACACCACATCGTAAAGTGAAAT

*TAS1c*

miR173a TS

syn-tasiR-Su

***>miR482aTS-NbSu***

GTGGTATGGGGGGAGTCGGGAATAGACCATTTATGTATGACTCCCGGAATTCCA

*TAS1c*

miR482 TS

syn-tasiR-Su

***>miR173TS-Su***

GTGATTTTTCTCTACAAGCGAATAGACCATTTATGTATGACTCCCGGAATTCCA

TAS1c

miR173 TS

syn-tasiR-NbSu

***>miR482TS-GUS(x4)***

GTGGTATGGGGGGAGTCGGGAATAGACCATTTATATTGACCCACACTTTGCCGATAACCTTCACCCGGTTGCCACTATTGACCCACACTTTGCCGATAACCTTCACCCGGTTGCCAC

*AtTAS1c*

NbmiR482a TS

syn-tasiR-GUS_Nb_-1

syn-tasiR-GUS_Nb-_2

***>miR482TS-TSWV(x4)***

GTGGTATGGGGGGAGTCGGGAATAGACCATTTATGTAAGACGTGATTGTGTCCTTATCAGCTCTGGGTGAATCGGTTGGTATAGTGGGGCATACCGTCAGAGTGCACAATCCATCTT

*AtTAS1c*

miR482 TS

syn-tasiR-TSWV-1

syn-tasiR-TSWV-2

syn-tasiR-TSWV-3

syn-tasiR-TSWV-4

***>miR173TS-TSWV(x4)***

GTGATTTTTCTCTACAAGCGAATAGACCATTTATGTAAGACGTGATTGTGTCCTTATCAGCTCTGGGTGAATCGGTTGGTATAGTGGGGCATACCGTCAGAGTGCACAATCCATCTT

*TAS1c*

miR173 TS

syn-tasiR-TSWV-1

syn-tasiR-TSWV-2

syn-tasiR-TSWV-3

syn-tasiR-TSWV-4
